# SISE, free LabView-based software for ion flux measurements

**DOI:** 10.1101/2025.04.09.647937

**Authors:** Namrah Ahmad, Krishani Tennakoon, Rainer Hedrich, Shouguang Huang, M. Rob G. Roelfsema

**Affiliations:** Molecular Plant Physiology and Biophysics, Julius-von-Sachs Institute for Biosciences, Biocenter, Würzburg University, Julius-von-Sachs-Platz 2, D-97082 Würzburg, Germany; Faculty of Synthetic Biology, Shenzhen University of Advanced Technology, Shenzhen 518055, China; Key Laboratory of Quantitative Synthetic Biology, Shenzhen Institute of Synthetic Biology, Shenzhen Institutes of Advanced Technology, Chinese Academy of Sciences, Shenzhen 518055, China

## Abstract

Plant growth and development strongly depend on the uptake of soil minerals and their distribution within plants. Various electrophysiological techniques have been developed to study these ion transport processes, from the single molecule-to whole plant level. An important non-invasive method is provided by Scanning Ion-Selective Electrodes (SISE), which are used to detect ion fluxes. These SISE-measurements depend on software that coordinates the perpetually electrode movement between two positions, as well as data collection and analysis. We developed two LabView-based programs; the SISE-monitor and SISE-analyser that enable ion flux recordings and their analysis, respectively. These applications are freely available, both as windows-executable files that enable routine measurements, as well as the LabView source code that allows deep insights into the routines used for measurement and further development of the programs to include new functions.

## Background

Ion transport plays a major role in the growth and development of plants. Nutrients are taken up from the soil in their aqueous ionic form and transported within the plant body, via xylem and phloem network. At their destination, these nutrients enter the cells of growing tissues, where they support osmotically driven cell expansion, as well as a multitude of specific functions. Because of the central role of ion transport in the physiology of plants (1), a variety of electrophysiological techniques have been developed to study these processes.

In the 1990^ies^, techniques were established to study plant ion fluxes with extracellular ion-selective electrodes. This minimal invasive technique enabled measurements of ion transport with intact tissues and single cells, such as leaves of aquatic plants (2), roots (3, 4) and pollen tubes (5, 6). For large tissues, such as leaves of the aquatic plant *Potamogeton lucens*, two miniature pH electrodes (diameter 1,5 mm) could be used at a fixed distance of 5 mm, to evaluate H^+^ fluxes in and out of leaves. These calculations of H^+^ fluxes were based on pH-gradients in an unstirred layer of solution, on the leaf surface (2).

Studies on smaller tissues, or single cells, required smaller ion-selective electrodes, which can be constructed from pulled glass capillaries, plugged with liquid membranes. The tip size of these electrodes can be as small as 10 µm, which enables measurements on single cells (5). However, the voltage of these electrodes tends to drift, which may due to loss of components from the ion-selective cocktail in the tips of these electrodes, or an imperfect seal between the cocktail and the silanized glass capillary (7, 8). Consequently, it is difficult to use two liquid membrane electrodes positioned at a fixed distance, as explained above for *P. lucens*. For this reason, a single ion-selective electrode is used, which moves perpetually between two positions. The use of such “self-referencing” scanning electrodes has become established and has been described in several method papers (9-12).

Even though all studies with self-referencing ion-selective electrodes are based on diffusion in unstirred layers of solution, different approaches have been developed to calculate the ion fluxes (11, 13). Whereas the group of Smith et al. (1994) uses Fick’s law of diffusion to determine the ion fluxes, Newman (2001) developed an approach that is based on the difference in chemical potential between two positions. At most conditions, both approaches will give similar results, but the method of Newman takes into account the impact of an electrical field on ion movement (Newman 2001) and thus will give better results if an electrical field is present in the bath solution.

Despite of using the same technical approach, the ion flux measurements methods have obtained various names, such as ion-selective vibrating-microelectrode system (4), scanning ion-selective electrode technique (SIET) (14), microelectrode ion flux estimation (MIFE) (12), or Non-invasive Micro-test System (NMT) (15). To some extend these names are protected and we therefore will use description Scanning Ion-Selective Electrode (SISE), or just ion flux measurement.

Central to ion flux measurements is a software that controls the electrode movement and collects data from the amplifier that is connected to the ion-selective electrode. In the period from 2013 to 2015, we encountered limitations with commercial software and started to develop two LabView-based programs to conduct, and analyse, ion flux measurements. LabView provides a graphical environment to program “test and measurement” software that is widely used by engineers and scientists. A community edition is provided that can be downloaded for free and used for non-commercial projects (https://www.ni.com/en/shop/labview/select-edition/labview-community-edition.html). The software is especially designed to support data acquisition and control measuring devices (https://www.ni.com/en/shop/labview.html) and thus very well suited to control electrode movements and data collection during ion flux measurements.

The SISE-Monitor offers a user-friendly approach to registers time-dependent changes in the voltage signal of the ion-selective electrode, which enables the experimenter to judge the validity of measurements. A second program, the SISE-Analyser, allows the re-evaluation of time-dependent voltage changes and calculate the ion fluxes, offline. These options were not available in the commercial software that we obtained in 2013 and have been crucial for ion flux measurements that were conducted in several research projects since 2018 (16-19). The LabView source code is now made publicly available through GitHub (https://github.com/Rob-Roelfsema/SISE-Software-April2025), which enables insights into the routines used in the SISE programs and allow new features to be implicated.

## Results

These SISE-software is separated in two applications; i) the SISE-Monitor that supports the measurements in real time and ii) the SISE-Analyser to conduct offline analysis and calculate ion-fluxes. Below the basic properties of both applications will be described. A point-by-point manual that explains how the software is installed and used, as well as a protocol to test the SISE measurement setup are provided in the supplementary materials.

### SISE-Monitor

Two versions of the SISE-Monitor will be provided as supplementary material and via GitHub. The SISE-Monitor100 is a fully active application that can be used in combination with the Patchstar micromanipulator (Scientifica, Uckfield, UK) and various analog-to-digital converters of NI (National Instruments Corp., Austin, TX, USA). In addition, a demonstration version is supplied (SISE-monitor100-demo), which provides an impression of the abilities of the program, without having to invest time and money into hardware components. Both programs are available as compiled programs (*.exe), and as “virtual instrument” (*.vi) files. The *.exe files run on a windows computer, in combination with a LabView runtime engine. In the supplementary files, version 2025 (Q1) of the runtime engine will be provided and later versions of the LabView runtime engine can be downloaded from the NI-website for free. The *.vi files can be opened in LabView and will allow modification of the program. The description below will apply to the normal SISE-Monitor100 program, but most of it is also valid for the demo version.

The monitor program consists of three main modules (Fig. 1), which are meant to be executed sequentially. In the first module, information needs to be entered about the hardware, data file and experimental conditions. The measurements are caried out in the second module and data storage is secured in the last module.

**Figure 1.**
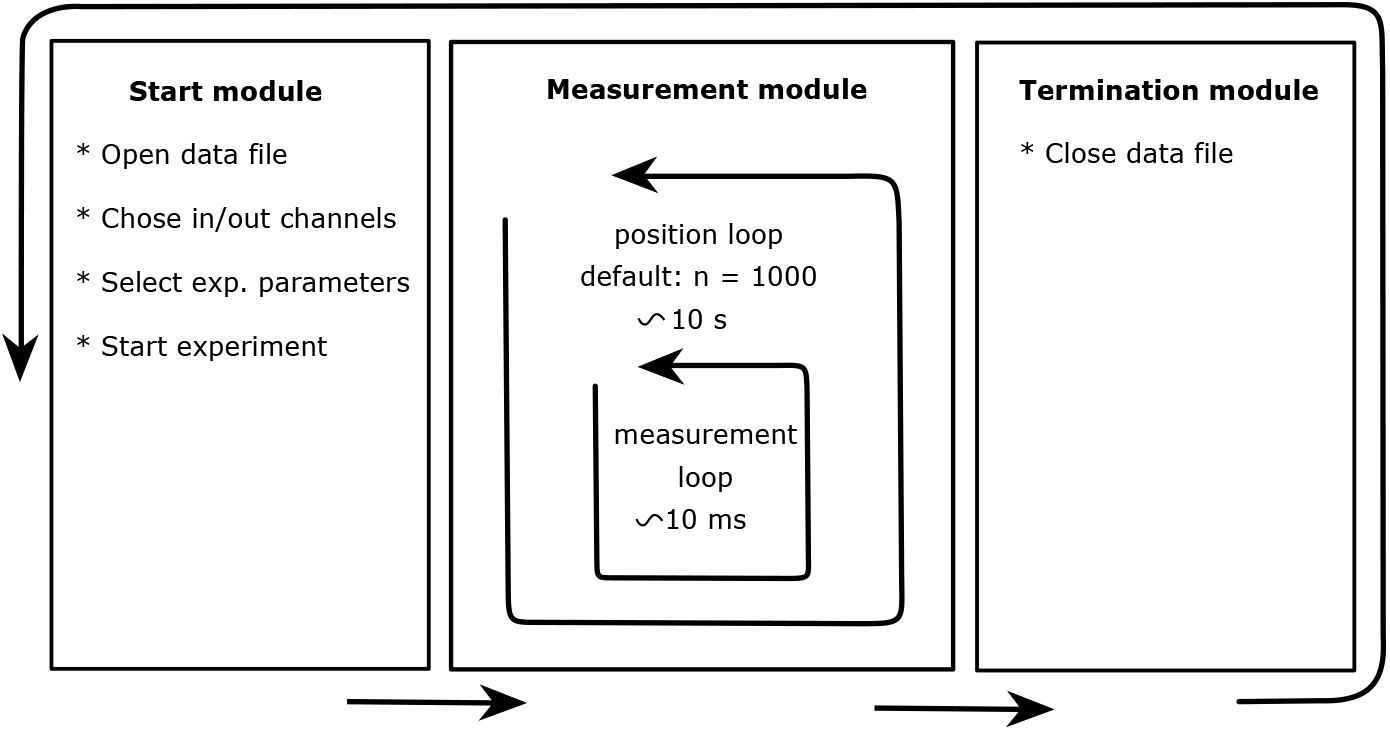
Schematic overview of the main modules of the SISE-Monitor. In the start module, a data file is generated, the input channels are chosen, and experimental parameters are selected, before the program is started. In the measurement module, a fast “measurement loop” is active that stores data at an interval of approximately 10 ms. At default conditions, the measurement loop is executed 1000 times within the “position loop” and thus adds up to a total time of approximately 10 s at one position. In the next “position loop” the electrode moves to the other position and again 1000 datapoints are acquired at default conditions. The data file is closed in the termination module and the next measurement can be started.

#### Start-up module

Before the SISE-monitor can be started, information needs to be provided about the hardware and experimental conditions (Fig. 2). First, the communication (COM) port of the Patchstar micromanipulator (Scientifica) has to be selected. If the correct port is chosen, the virtual LED next to the COM-port window, will turn green. Below the COM-port window, the speed of the manipulator can be changed from a fast mode, which is useful to position the electrode, to the slow mode that is required to start the SISE recording. Next, four input channels of the analog-to-digital (A/D) converter need to be selected (refer to the manual for further details) and a data file name must be provided.

**Figure 2.**
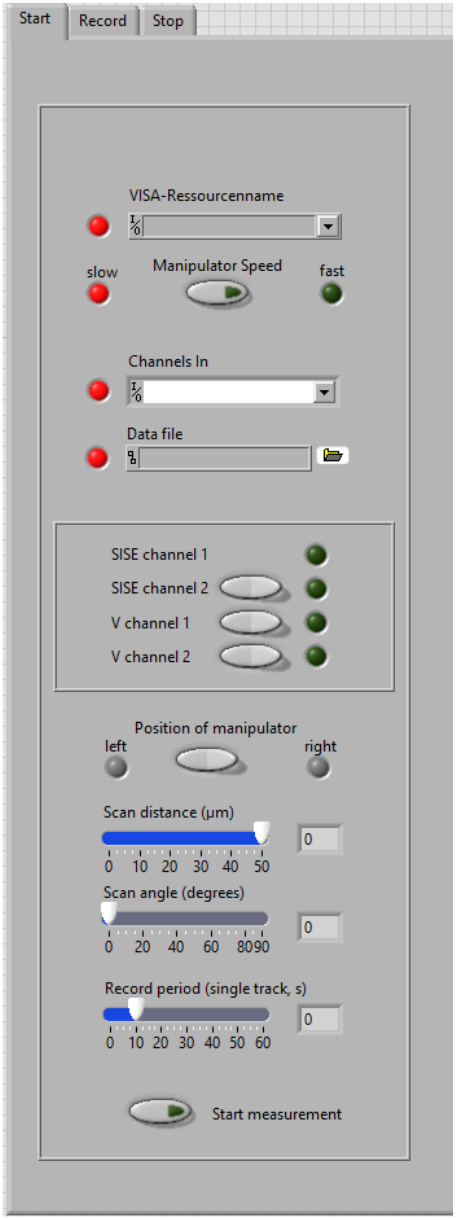
Start window of the SISE-monitor. *Upper part:* Before the SISE-monitor can be started several parameters have to be entered. In the upper “VISA-Resource name” window, the COM-port must be chosen to which micromanipulator is connected. The virtual LED next to the window will turn green, once a parameter is correctly entered. The manipulator can be switched between a fast and slow mode with the “Manipulator Speed” switch. The measurement can only be started with the manipulator in the slow speed mode. In the “Channel in” window, four channels of the NI-interface need to be selected, which correspond to the SISE channel 1 and 2 and Voltage channel 1 and 2, respectively. In the window below, a data file name must be provided. *Middle part*. The switches allow selection of the data channels that will be recorded in addition to SISE channel 1. *Lower part:* A switch is provided to enable measurements with a manipulator positioned on the right, or left side, of the tissue of interest. In addition, three sliders are available to set the scan distance of the ion-selective electrode, the angle at which the electrode moves (0 degrees is horizontal) and the period of measurement at one position. The measurement can be started, once all virtual LEDs are green, by pushing the switch at the bottom of the window.

The program can be used with a variety of NI A/D converters, but it is advisable to use high-end A/D converters, to minimize the noise level and thereby enhance the resolution of the measurements. The first chosen analog input channel (in most cases AI0) should be connected to the first ion-selective electrode amplifier. A second SISE channel can be selected, by pushing the virtual switch (“SISE channel 2”), just as two additional voltage channels. These data source channels will be assigned as; AI1, AI2 and AI3, to further analog input channels. In total the program thus can store data of 4 channels; 2 SISE channels and 2 additional voltage channels.

Measurements can be conducted on the left, or right side, of the tissue that is studied. The position of the manipulator has to be provided, to ensure that first movement of the electrode is away from the tissue (or cell). Finally, 3 parameters need to be selected that define the distance, the angle, and the interval by which the micromanipulator moves. Note that the selected record period may slightly differ from the actual period at which the system measures at a certain position. The program can be slowed down by other processes that are running on the computer, which elongate the recording time. However, the exact recording time will be stored in the data file.

Once all virtual LEDs have turned green, the measurements can be started.

#### Measurement module

The movements of the micromanipulator are controlled by “Virtual Instrument Software Architecture” (VISA) comments, which can be downloaded, as a NI-VISA file, from the NI website (https://www.ni.com). Before the start of the experiment, the electrode tip must be moved manually to position 0, close to the tissue of interest (the distance should be documented). Upon start of the experiment, the electrode will remain at position 0 during the first measurement and only will move away from the tissue, to position 1, during the next measurement (fig. 3).

**Figure 3.**
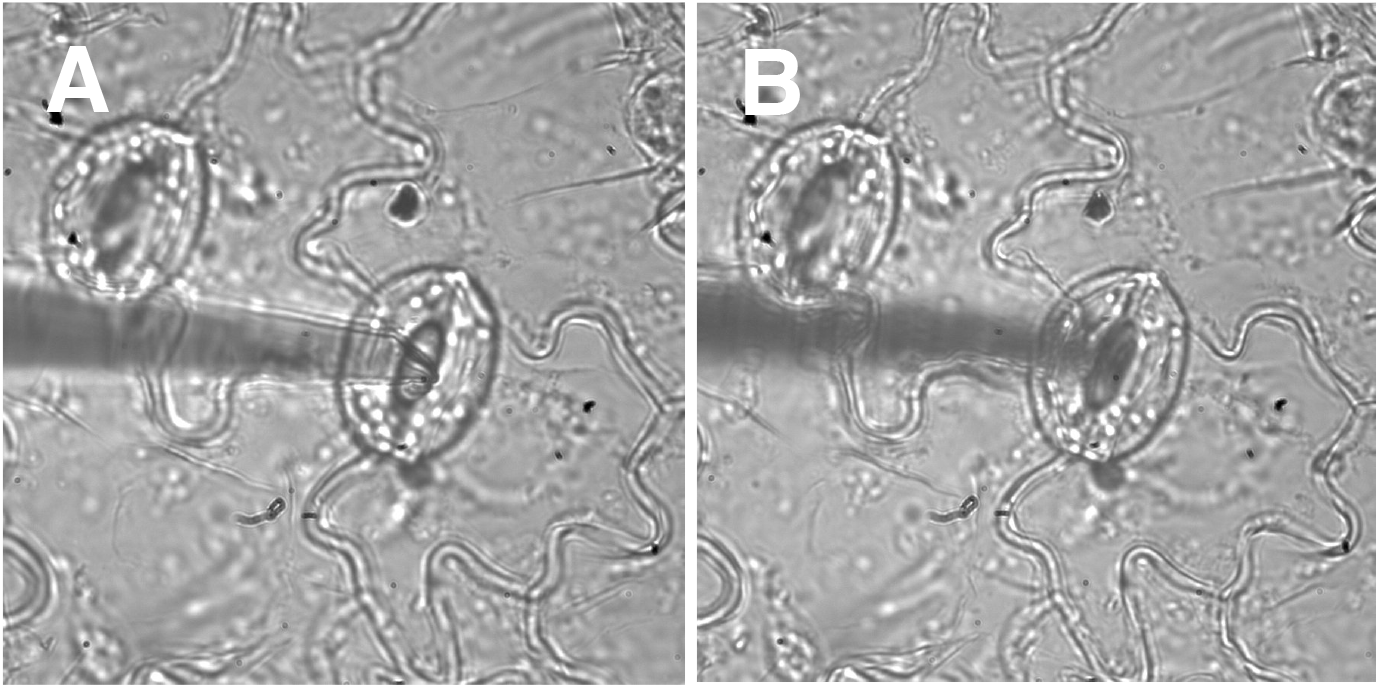
Ion flux measurement on an abaxial epidermal strip of a *Vicia faba* leaf. **A**. An H^+^-selective electrode with its tip at position 0, close to the stoma of interest. **B**. The tip of the electrode moved 50 µm upwards, to position 1. Note that the tip of the electrode moved out of focus.

By default, the electrodes move horizontally (angle of 0°), over a distance of 50 µm, every 10 s, but these values can be altered before the start of the experiment, as explained above. In experiments with epidermal strips of *Vicia faba*, we study guard cells with ion-selective electrodes that move 50 µm vertically (angle = 90°). Such a measurement is shown in Fig. 3, in which the electrode is in focus in position 0 (Fig. 3A, close to the stoma), while its tip moves out of focus to position 1 in Fig. 3B. In this measurement, an H^+^-selective electrode was used to study H^+^-fluxes induced by application of NO_3_^-^, as shown in Fig. 5 and explained for the SISE-Analyser below.

#### SISE channel data collection

During the measurements, the position of the scanning electrode is registered every 10 ms and displayed in the upper middle graph of the measurement window (Fig. 4). The concurring change in the voltage signal is documented in central graph, while the lower middle graph depicts the procedure to determine the voltage difference between position 0 and position 1.

**Figure 4.**
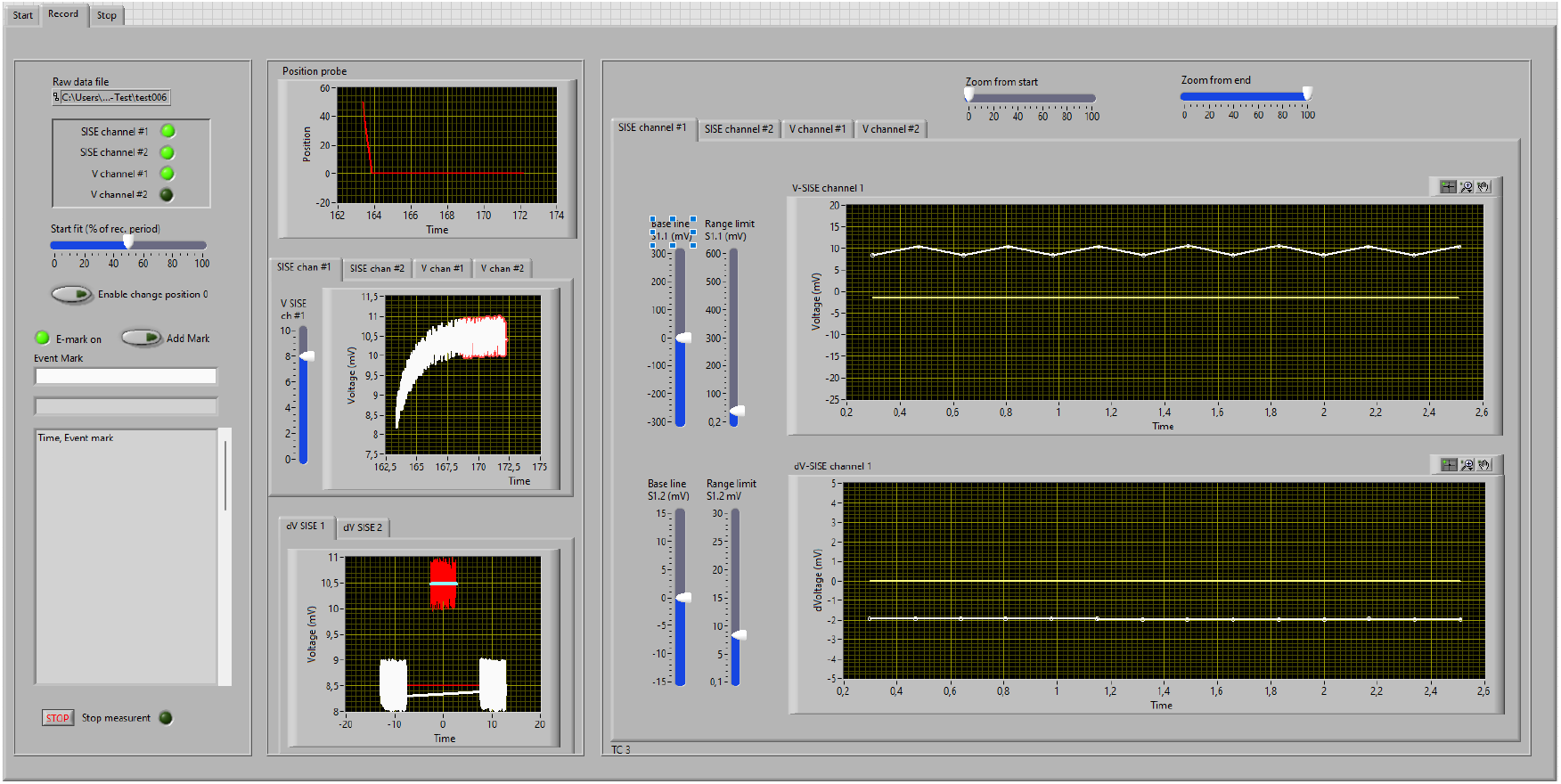
Measurement window of the SISE-Monitor (demo Version). *Windows on left:* Upper window shows overview of the channels that are recorded. The “Start fit” slider allows selection of the period of the measurement that is used for calculation of the voltage difference between position 0 and position 1. The lower windows can be used to enter “event marks” that will be stored in the data file. *Windows in the middle:* Upper graph; the position of the electrode in µm distance from position 0 plotted against time. Middle graph; the voltage signal of the electrode, plotted against time (in the demo version a slider is provided to manipulate the voltage). Lower graph; the voltage difference between position 0 and 1 is determined, by comparing the linear regression of two measurements at one position with that of the other position. Only the last part of the voltage recordings (set with the “Start fit” slider) is taken into account. *Windows on right:* Voltage signal (upper graph) and the voltage difference between position 0 and 1 (lower graph) plotted against time (in min.). The base line and voltage range can be changed with sliders on the left, while the slider above the graph can be used to alter the time range.

The voltage signal of the ion-selective electrode is measured with a microelectrode amplifier (VF-102, Bio-Logic, Claix, France, or similar). This amplifier is equipped with a high impedance headstage (input impedance > 10^14^ Ω, HS111, Bio-Logic, or similar), while a custom-made low pass filter, which also amplifies the signal -33 times, is mounted to the output (see also Test Protocol, Fig. 1). The minimal noise level of this system is 30 µV, using the NI-USB-6259 A/D converter. However, depending on the properties of the ion-selective electrode, it is likely that the noise level in normal SISE-recordings will be considerably higher.

Voltage data acquired by the amplifier are plotted against time in central graph of the measurement window (Fig. 4). Due to the slow exponential response of ion-selective electrodes the voltage signal will be delayed, compared to movement of the electrode. By default, the voltage of the electrode is assumed to become stable after 5 min., this period is indicated by a “red shadow” behind the white curve, and these data points are used to calculate the average voltage values and voltage difference between position 0 and position 1. Depending on the response time of the electrodes, the experimenter can adjust the period in which data are used for further analysis with the “start fit” slider during the measurements. Note that the position of the “start fit” slider does not affect the stored data and the period at which the data are averaged can be modified offline in the SISE-Analyser.

Because of the tendency ion-selective electrodes to float with respect to the voltage signal, a procedure was developed Newman (2001) to correct voltage drift. This procedure is depicted in the lower middle graph of the measurement window (Fig. 4). It shows the selected data points (starting at the time point chosen with the start fir slider) of the last three measurements. The first and last measurements are at the same position (position 0 in Fig. 4), while data in the middle were collected at the other position. Next, a regression line is obtained for the data points of the first and last measurement, followed by calculation of the regression line through datapoints of the middle measurement, using the same angle as found for the first regression line. The dV (V_position0_-V_Position1_) value can be deduced from the distance between both regression lines (2 mV in Fig. 4). These dV values are plotted against time in the right window, lower graph (Fig. 4) and indicate the changes in ion fluxes in time.

#### Data storage

Data are stored in an American Standard Code for Information Interchange (ASCII)-text file during each cycle of the measurement loop (approximately 10 ms). Comments to the experiment can be entered in the “Event Mark” window and will be stored in the “Event Mark” list and datafile, after clicking the “Add Mark” button. This list will also show, if the position of the manipulator is changed during the measurements, which can be enabled by activating the “Enable Change Position” switch (Fig. 4, left panel). The measurement is stopped by pushing the “stop button” at the bottom of the left window (Fig. 4). Once the “stop button” is activated, the last position loop will be completed, followed by the activation of the “termination module” (Fig 1). In the termination module the data file is closed, and a new measurement can be started by selecting “New measurement”, or the application can be stopped with “Stop program”.

### SISE-Analyser

#### Loading and scrolling data

Data obtained with the SISE-Monitor can be viewed with the SISE-analyser and converted into ion concentrations and ion-flux rates. In the first step, a data file is re-analysed using the same procedures as in the SISE-Monitor, to average data and calculate the voltage difference between position 0 and position 1. To this purpose, scroll switches can be used to view each measurement and recalculate the average voltage data (V) and voltage difference signals (dV) (Fig. 5). If all measurements have been covered, the virtual LED next to the scroll switches will turn green and allow the determination of ion concentrations and fluxes.

**Figure 5.**
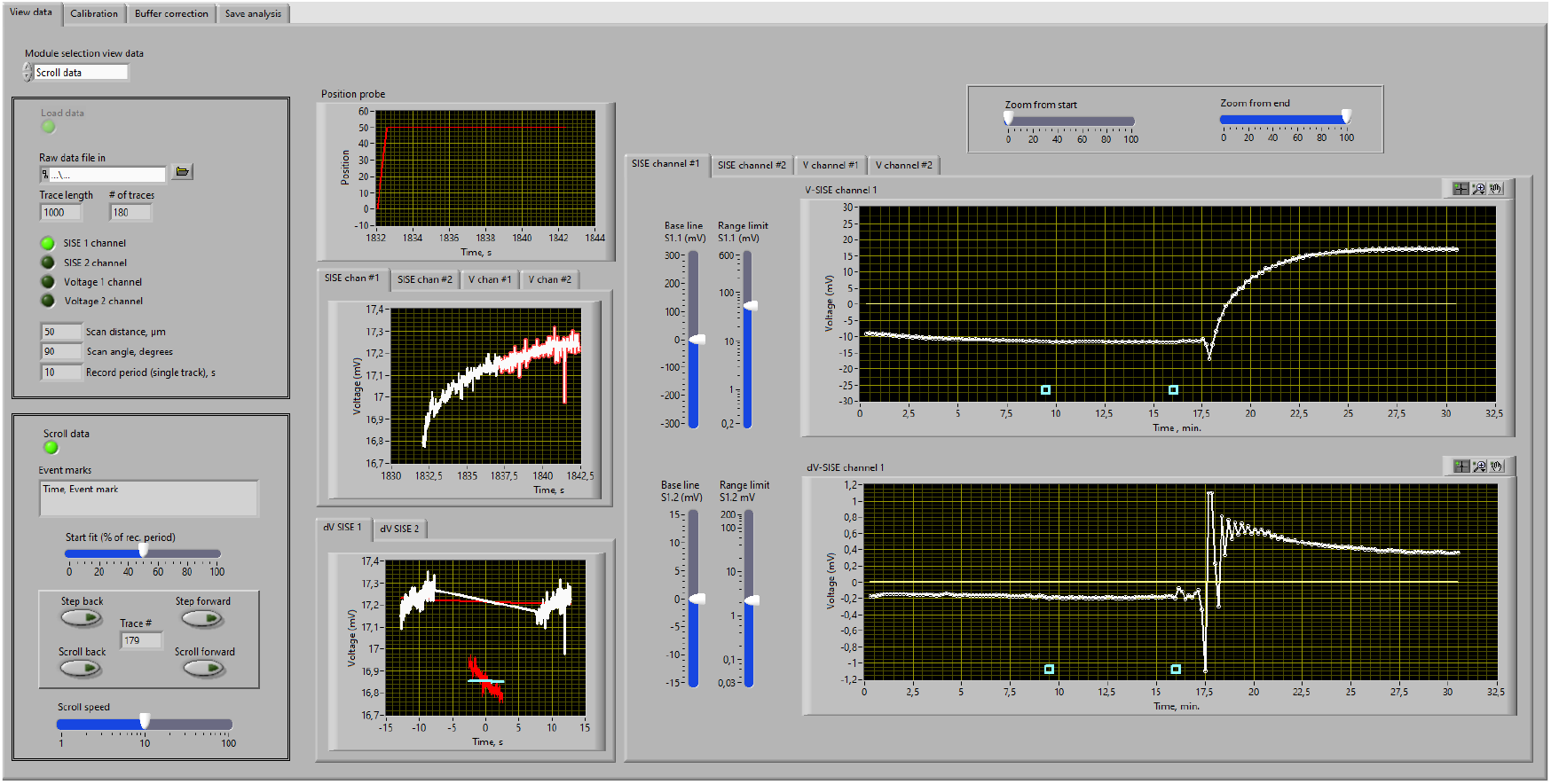
View data window of the SISE-Analyser. *Windows on left:* Sub-modules within the “View data window” can be activated with the selector in upper left corner. A datafile is uploaded and parameters of the measurement are displayed in the “load data” module. The data can be viewed in the “Scroll data” window, in which “step back” and “step forward” enable to switch between measurements and the “Scroll back” and “Scroll forward” allow rapid scrolling of the data. *Windows in the middle:* Upper graph; the position of the electrode in µm distance from position 0, plotted against time (s). Middle graph; the voltage signal of the electrode in time (the measurement shown was conducted with a pH-electrode close to guard cells at position 0 and 50 µm above the stoma at position 1). Lower graph; the voltage difference between position 0 and 1, determined by comparing the linear regression of two measurements at one position with that of the other position. Only the part of the voltage recording is used, which is indicated by the “red shadow” in the central graph (set with “Start fit” slider). *Windows on right:* Voltage signal (upper graph) and the voltage difference between position 0 and 1 (lower graph). The base line and voltage range can be changed with slider on the right, while the slider above the graph can be used to alter the time range.

In Fig. 5 the analysis of a H^+^-flux measurement on a stoma of *V. faba* is shown. At the start of the experiment a small efflux of H^+^ is evident from the negative dV value (Fig. 5, lower right graph). The addition of 10 mM LiNO_3_^-^ after 16 min. caused an increase of the voltage measured by pH electrode (Fig. 5 upper right graph) and a change of dV to positive values that indicates the uptake of H^+^ into guard cells (Fig. 5 lower right graph).

#### Calculation of ion fluxes

Ion fluxes are determined with the aid of the chemical potential difference of ions at position 0 and position 1. These calculations are carried out in the “Calibration” module (Fig. 6), in which several sub-modules are included. Upon selection of the “Ion species” sub-module, the ion species of interest can be chosen, and the ion mobility is displayed (data were obtained from the literature as explained in the supplement part 3, ion mobility data). The “Calibration” sub-module is used to determine the calibration curve (left window, Fig. 6). To this purpose, values must be entered for the ion concentrations in the calibration solutions (in mM) together with corresponding voltage signals (in mV). As soon as these values are entered, a calibration graph will appear in the window of this sub-module and the virtual LED will turn green.

**Figure 6.**
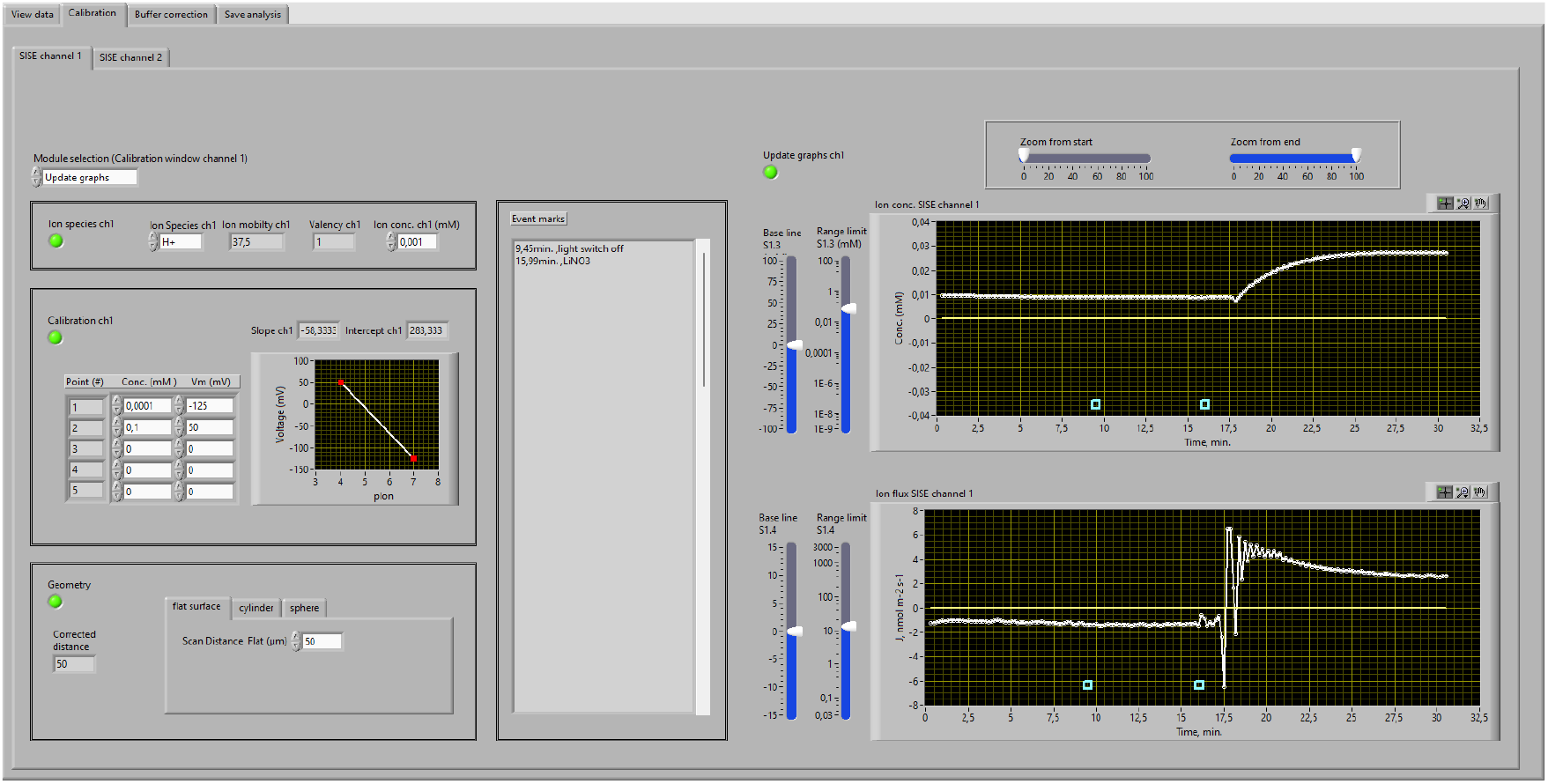
Calibration window of the SISE-analyser. *Windows on left:* Sub-modules within the “Calibration window” can be activated with the selector in upper left corner. In the “Ion species” module, the left window enables the selection of the proper ion species and its concentration in mM needs to be entered in the right window. Calibration values of the ion-selective electrode are entered in the table of the “calibration” window and are shown in a graph. The size and shape of the tissue, or cell, needs to be provided in the “Geometry module”, in which a corrected distance is calculated for objects with a cylindrical, or spherical shape. *Middle window:* The “event marks” that were entered during the experiment will be shown in the “Event marks” list. *Windows on right:* Ion concentration in mM (upper graph) and the calculated ion flux (H^+^ flux in this experiment) (lower graph) plotted against time (in min.). The base line and voltage range can be changed with slider on the right, while the slider above the graph can be used to alter the time range.

In the experiment shown in Fig. 6, the addition of 10 mM LiNO_3_^-^ caused acidification of the bath solution (upper right graph), while the H^+^ flux changed from approximately −1.5 to 2.6 nmol m^-2^ s^-1^ (lower right graph).

In case that the measured tissue can be regarded as a flat surface, only the distance between position 0 and position 1 needs to be entered (in µm), in the “Geometry” sub-module. Corrections for cylindrically shaped tissues, or spheres, are made as described by Newman (2001). The “Corrected distance” of movement is shown on the left side of the “Geometry” window. Once all information of the sub-modules (left windows in Fig. 5) is available (virtual LEDs turn green), the “update graphs” window can be selected and ion concentrations are calculated (Fig. 6, right window, upper graph). The ion fluxes (Fig. 6, right window, lower graph) are determined with the use of the equation 01, derived by Newman (2001) (11).

Equation 01,

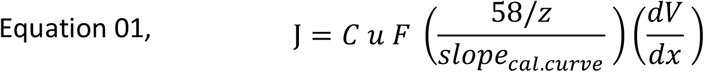

in which J is the ion flux (in mol m^-2^ s^-1^), C is the average concentration of the ion (in mol m^-3^), u is the mobility of the ion (in (m s^-1^)(N mol^-1^)^-1^), 58/z the slope of an ideally ion-selective electrode divided by the valency of the ion, slope_cal.curve_ is the actual slope of the calibration curve, dV, the voltage difference between position 0 and position 1 and dx the corrected distance of movement of the electrode.

#### Buffer correction

The calculated ion flux data will resemble the flux of free ions. However, if the ion is buffered in the solution, it will form a complex with the buffer molecules and cause a gradient of conjugated buffer molecules. In other words, part of the H^+^ flux across the plasma membrane will cause a flux of conjugated buffer, instead of a flux of free H^+^. For this reason, a correction procedure for pH buffers has been developed by Arif et al. (1995) that is integrated in the SISE-analyser “Buffer correction” module. Within this module, common buffers such as 2-(N-morpholino) ethanesulfonic acid (Mes) and tris(hydroxymethyl)aminomethane (TRIS) can be selected (for mobility see supplement part 3). In addition, some organic acid buffers are provided but note that these buffers can have several pK_S_ values and the values only can only be used if the pH of the solution is close (less then 0.25 pH units difference) to the pK_S_ value that is indicated. Additional buffers can be introduced in the VI-version of SISE-analyser using the community edition of LabView.

The ion flux values are corrected for the flux of the protonated buffer, based on the following equation (Arif et al., 1995):

Equation 02,

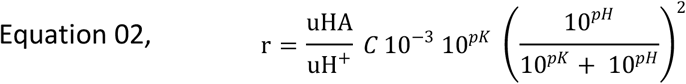

where r is the ratio between the flux of H^+^ and the protonated buffer (HA), u_HA_ the mobility of the protonated buffer, u_H+_ the mobility of H^+^ and C is the total concentration of the buffer ([AH] + [A^-^]). The total flux is calculated as: J_total_ = (r+1) J_H+_.

## Data storage

Storage of the analysis occurs in the “save analysis” module, in which the “analysis output file” is entered and data can be selected, which will be written in a text file. Depending on the number of channels that were activated during the measurement, and the type of analysis (with, or without buffer correction), the “analysis out file” can have up to 14 columns of data.

## Discussion/Conclusion

The SISE-programs that are provided in the supplementary data files, and on GitHub, will enable routine ion flux measurements and the analysis of these recordings. Because of the option to download the source code (virtual instrument in LabView), the application can be easily updated and adapted to future versions of Microsoft Windows.

There are several additional features that we intend to introduce into the SISE-programs. One of these features is to capture images that document the position of the electrode, relative to the object that is studied. This option was already implemented by Shipley and Feijo (1999) and is especially useful, if ion fluxes are determined at range of positions along the surface of a tissue. Such an experimental approach has provided data of ion-uptake gradients, along pollen tubes (5, 9, 20), or roots (21, 22). Image capturing and analysis thus will be a valuable extension planned to be included in future versions of the SISE-applications.

Camera control and image analysis can be conducted with “Micromanager”, a free software package that enables simultaneous control of microscope cameras and illumination devices (23). The combined use Micromanager and the SISE-Monitor will enable imaging and ion-flux measurements simultaneously. Moreover, the Micromanager can be used to conduct measurements with genetically-encoded fluorescent ion-indicators. The simultaneous used of Micromanger and the SISE-Monitor application thus may enable linking extracellular ion fluxes to changes in the cytosolic ion concentration of plant cells.

Micromanager uses the ImageJ application (24) to visualize and analyse the images that are captured. This program thus will also be useful to analyse the captured images, in combination with ion flux analysis that is carried with the SISE-Analyser. We intend to synchronize the ion flux- and image data-analysis and thus enable a rapid and easy procedure to link the results of both methods.

Because of the availability of the source code in the virtual instrument-version of the SISE-programs, it can be modified and expanded by any of the users. We would welcome, if such new attributes of the SISE-programs would be made publicly available. This could provide a range of versions of the SISE-programs, with a variety of helpful features that would enable SISE-users to find an optimal solution for their needs.

## Availability and requirements

Project name: SISE-Software

Project home page: https://github.com/Rob-Roelfsema/SISE-Software-April2025

Operating system: Windows

Programming language: LabView

Other requirements: Supporting Virtual Instrument (VI) files (for *.vi files only)

License: GNU GPL

Any restrictions to use by non-academics: none

## List of abbreviations

AI channel: Analog Input channel
ASCII: American Standard Code for Information Interchange
A/D converter: analog-to-digital converter
MIFE: Microelectrode Ion-Flux Estimation
NI: National Instruments Corp.
NMT: Non-invasive Micro-test system
SIET: Scanning Ion-selective Electrode Technique
SISE: Scanning Ion-Selective Electrode
VISA: Virtual Instrument Software Architecture
^*^.exe: Windows executable file
^*^.vi: LabView virtual instrument file.

## Declarations

## Ethics approval and consent to participate

Not applicable

## Consent for publication

Not applicable

## Availability of data and materials

The software will be available in the supplementary materials and on GitHub, as *.exe and *.vi files.

## Authors’ contributions

NA, KT and SH, tested the software and contributed to writing of the manuscript, RH initiated the project and contributed to writing of the manuscript, MRGR initiated the project, wrote the software and contributed to writing of the manuscript.

## Acknowledgements

We thank Ulrich Schliwa, Ulrich Turba, Michael Dienesch and Thomas Walter (University of Würzburg, Germany) for support and advice with electronics.

## Supplementary files

1. Manual for the SISE-Monitor and SISE-Analyser
2. Test protocol for the SISE setup.
3. Table with ion mobility values

