## Supplemental information for "SISE, free LabView-based software for ion flux measurements"

**Contents:** **Pages**


1. SISE-Monitor and Analyser manual 2-11
2. Protocol for testing the SISE-setup 12-15
3. Ion mobility data 16-17

### SISE-Monitor and Analyser manual, version 1 (2025)

### Content:


| Content: | Page |
|---|---|
| **Software installation** | **3** |
| **SISE-Monitor and SISE-Monitor-demo** | **4** |
| A. Initiation and start of the program | 4 |
| A1. VISA COM-port for Patchstar manipulator | 4 |
| A2. Input channels of the analog/digital converter | 4 |
| A3. Data file entry | 4 |
| A4. Selection of input channels | 4 |
| A5. Right/left site manipulator | 5 |
| A6. Select the distance, scan angle and record period | 5 |
| A7. Start of measurement | 5 |
| B. Record module | 5 |
| B1. Pilot window of record module | 5 |
| B2. Graphs of voltage (V) and differential voltage (dV) values | 6 |
| B3. Graphs of time dependent changes in V and dV | 6 |
| C. Stop module | 7 |
| **SISE-Analyser** | **7** |
| D. Start of the application and data browsing | 8 |
| D1. Load data file | 8 |
| D2. View measurements in data file | 8 |
| E. Calculation of ion concentrations and fluxes | 9 |
| E1. Selection of ion species | 9 |
| E2. Calibration of ion selective electrode | 9 |
| E3. Geometry of tissue, or cell | 9 |
| E4. Graphs of calculated ion concentrations and fluxes | 10 |
| F. Buffer correction | 10 |
| F1. Buffer species | 10 |
| F2. Concentration of buffer | 10 |
| F3. Graphs of buffer-compensated ion fluxes | 10 |
| G. Save analysis | 10 |
| G1. Create analysis file | 10 |
| G2. Select channels for storage | 11 |
| G3. Start new analysis/terminate program | 11 |
| **References manual** | **11** |

### Software installation

The SISE-programs are provided as Windows-executable (*.exe) files and NI (National Instruments) LabView virtual instrument files (*.vi). The SISE-Monitor100.exe and SISE-Analyser100.exe files require installation of the NI-LabView Runtime program (version 2025 Q1, Tab. 1) that can be downloaded from the NI -Website (https://www.ni.com/). The virtual instrument files only can be opened with a version of NI-LabView (Tab. 2), such as the community edition that is free for non-commercial use. Please follow the instruction of the NI-website to install either the LabView Runtime program, or a version of LabView.

In case that the virtual instrument version is used, several **sub-VI's** need to be assessable to the main VI application. These sub-VI's are stored in the folder "SubVIs-SISE-Analyser100", which is available in the ZIP-files that are provided in the supplementary materials and on GitHub (https://github.com).

Two versions of the SISE-Monitor are available, the SISE-Monitor-demo can be used to get an impression of the application, without the need to obtain an analog/digital (A/D)-converter and Patchstar manipulator. The fully functional SISE-Monitor application requires installation of the correct drivers for the A/D converter, which is provided by the NI-DAQ$^{TM}$ mx file. Moreover, the NI-VISA system must be installed to enable communication with the Patchstar (Scientifica, Uckfield, UK). Please follow the instructions of the NI-website to install NI-DAQ$^{TM}$ and NI-VISA.

Table S1, software components required for running the SISE "*.exe" applications.

| | SISE-Monitor-demo | SISE-Monitor | SISE-Analyser |
|---|---|---|---|
| SISE-application (*.exe) | + | + | + |
| NI-Runtime 2025 Q1 | + | + | + |
| NI-DAQ$^{TM}$ mx | - | + | - |
| NI-VISA | - | + | - |

Table S2, software components required for running the SISE "*.exe" applications.

| | SISE-Monitor-demo | SISE-Monitor | SISE-Analyser |
|---|---|---|---|
| SISE-virtual instrument (*.vi) | + | + | + |
| NI-LabView | + | + | + |
| NI-DAQ$^{TM}$ mx | - | + | - |
| NI-VISA | - | + | - |
| SISE sub-VI's | + | + | + |

If the proper software components are installed, the *.exe application can be opened in Windows, whereas the *.vi files are opened in LabView.

### SISE-Monitor and SISE-Monitor-demo

All steps will be described for the SISE-Monitor but will be similar for the SISE-Monitor-demo version. Differences between both applications will be indicated.

Note that the SISE-Monitor does not calculate ion concentrations and fluxes, but instead displays voltage- and differential voltage values, which can be used to estimate changes in the concentration and fluxes of the ion that is studied. The values of the ion concertation and flux values are determined in a later stage with the SISE-Analyser. However, the **ion-selective electrode must be calibrated** to support later analysis with the SISE-Analyser. A maximum of 5 calibration points can be entered in the SISE-Analyser.

The SISE-Monitor is started by clicking the white arrow 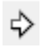 in the upper left corner of the screen. Once activated, the program can be stopped at any time by pressing the red button 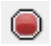.

**A. Initiation and start of the application.**

A1. Choose the VISA COM-port to which the control board of the Patchstar manipulator is connected, via a USB cable. If the correct channel is chosen, the virtual LED next to the "VISA-Resource name" window will turn green.
The speed of the manipulator can be switched between a slow and fast mode with the "Manipulator Speed" switch. The fast mode is convenient to position the electrode, while the SISE-measurement can only be started with the manipulator in the slow mode.

A2. Choose four input channels of the A/D-converter in the "Channels In" window, by clicking the arrowhead and select "Browse…" (Fig. 1A), in the next window select three channels, like channel ai0 to ai3 (Fig. 1B).

A

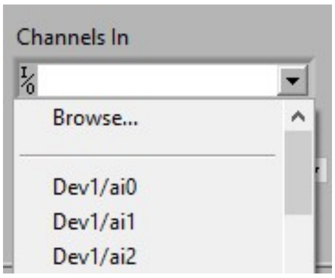


Figure 1A, 1th channel selection window

B

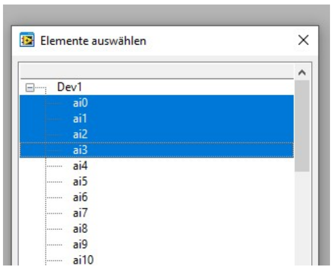


Figure 1B, 2nd selection window

Figure 1, Selection of the input channels of the analog/digital converter. **A.** In the Channels in window, click "Browse…" **B.** Select four AI channels and press enter.

A3. Enter a data file name in the "Data file" window. Only a none-existing file name will be accepted, the program cannot overwrite old data files.

A4. Choose the input channels of which data should be stored. The data of SISE channel 1 are stored by default, an additional SISE channel 2, and 2 additional voltage channels can be selected with the switches.

A5. Select if the manipulator is positioned on the left, or right side of the tissue/cell that is studied (the first movement will always be away from the tissue/cell).

A6. Select the distance, scan angle and record period, with three sliders in the lower window in the left.

a. Use the slider "Scan distance" to set the distance by which the electrode will move (position 0 to position 1) in µm, the default value of 50 µm can be changed in steps of 10 µm.

b. Select the angle by which the electrode will move with the slider "Scan angle" in steps of 5°. The electrode will move horizontally at 0° and vertically at 90°.

c. Use the "record period" slider to set the period of measurement at one of both positions. The default value is 10s, which can be changed in steps of 5s. Note that the real time of measurement may slightly deviate from the period selected period. The correct timing will be shown in the graphs of the "Records" window and stored in the data file.

A7. In case that all virtual LEDS's are green on the left side of the window, the measurement can be started by clicking the start measurement switch.

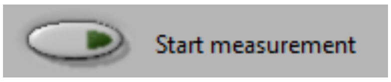


**B. Record module.**

*B1. Left window.* After the start of the recording, the "Record" window will automatically appear and display the following items in the left window:

a. The upper window indicates the data file to which the data are stored.

b. Active channels are indicated by green virtual LEDs in the second window.

c. The "Start fit" slider can be adjusted to change the time point during each measurement, from which data are used to calculate the voltage and differential voltage (dV) data. Note that this slider only affects the online analysis but not stored data.

d. The electrode movement is fixed between 2 positions, as long as the "Enable change position 0" switch is off. Switching the "Enable change position 0" on (virtual LED turns green) will allow moving the electrode to a new position. However, this option is only active at position 0, and not at position 1. Movement at position 1 is disabled to prevent movement of the electrode into the tissue (or cell) when it returns to position 0.
Changes in position are automatically recorded in the "Event mark list" in the lower window, together with the time at which the position was changed. This event mark list is stored in the data file.

e. Changes in experimental conditions can be entered in the upper "Event Mark" window. Press the "Add Mark" switch to mark the time at which experimental conditions were changed, the text will move into the next window. After a change in position of the electrode, the text and the time at which the "Add mark" switch was pressed, will appear in the lower window and stored into the data file.

f. The measurement can be stopped by pressing the STOP button. The last measurement will be finished, and the program will move to the "Stop" module.

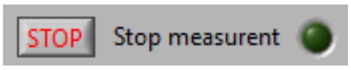

*B2. Middle window.* In the middle window three graphs show the following data:

a. The upper graph indicates the position of the electrode. Position 0 has the value of 0, while the movement to position 1 is shown in µm and values on the X-axis indicate the time in seconds. Note that during the first measurement the electrode remains at position 0, the first movement to position 1 occurs during the second measurement.
b. Voltage data of all 4 channels are plotted against time in the central graph. Each of the channels can be selected by clicking one of the following headers;

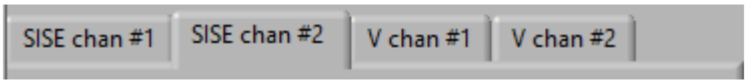


Part of the voltage curve is marked with a "red shadow". This period of the measurement is used for calculating the average voltage values and to determine the differential voltage between the two positions, as depicted in the lower graph. The "Start fit" slider can be used to alter this period.
In the demo version of the SISE-monitor, data are generated with a single exponential function, to which a "noise signal" is added. The magnitude of the voltage signal can be changed with the slider on the left side of the graph.
c. In the lower graph, the procedure is shown through which the voltage difference (dV) is determined between position 0 and position 1. The procedure uses three data sets, to reduce the impact of electrode drift on calculation of dV. If three measurements have been conducted, the graph shows voltage data plotted against time (s), of which the start is selected with the "Start fit" slider. The voltage difference between position 0 and 1 is calculated as the distance along the Y-axis between two regression lines. The first regression line is calculated with the first and last data set, while a second regression line has the same angle as the first one and is determined for the middle data set. Graphs can be selected for the SISE channel 1, or SISE channel 2.

B3. Right window. Time dependent changes in voltage (upper graph) and voltage difference (dV) between position 0 and 1 (lower graph) for each of the SISE channels are show. In addition, the data of voltage channels can be viewed by clicking the headers;

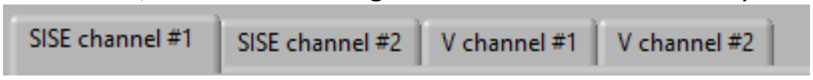


a. If one of the SISE channels is selected, the upper graph shows the voltage signal recorded by the ion selective electrode, plotted against time (min.). These data indicate changes in ion concentration, but actual ion concentrations are calculated with the SISE-Analyser. Note that the first data point is shown with a delay, only after three measurements have been conducted. This first data point shows the second measurement (at position 1).
The time window that is shown in the graph can be altered with the sliders "Zoom form start" and "Zoom from end" above the graphs. This will apply to all graphs in the right window. The scale of the Y-axis can be changed with the "Base line" and Range limit" sliders on the left side of each graph. Note that the time point of "Event marks" is indicated by a light blue square on the bottom of the graph.

b. The lower graph is only available for the SISE channels and shows differential voltage values, plotted against time (min.). These values indicate the magnitude of ion fluxes, while the SISE-Analyser is used to determine the ion flux values (nmol $m^{-2}$ $s^{-1}$). The differential voltage values are determined by the procedure described above for the central graph (point B2b). Because of the requirement of 3 measurements for this procedure, the first data point is shown for the second measurement (at position 1), after a delay. The range of the X-axis and Y-axis can be changed as explained for the voltage graphs above (point B3a).

#### C. Stop module

Only two options are provided in this module, either a new measurement can be started by clicking the “OK” button next to the “New measurement” window, or the program can be stopped by clicking the “STOPP” button. In both cases the data file will be closed and can be analysed with the SISE-Analyser application.

### SISE-Analyser

In the SISE-Monitor, data are stored as tab-separated American Standard Code for Information Interchange (ASCII) codes, which can be viewed in the editor of Windows, as shown in figure 2.

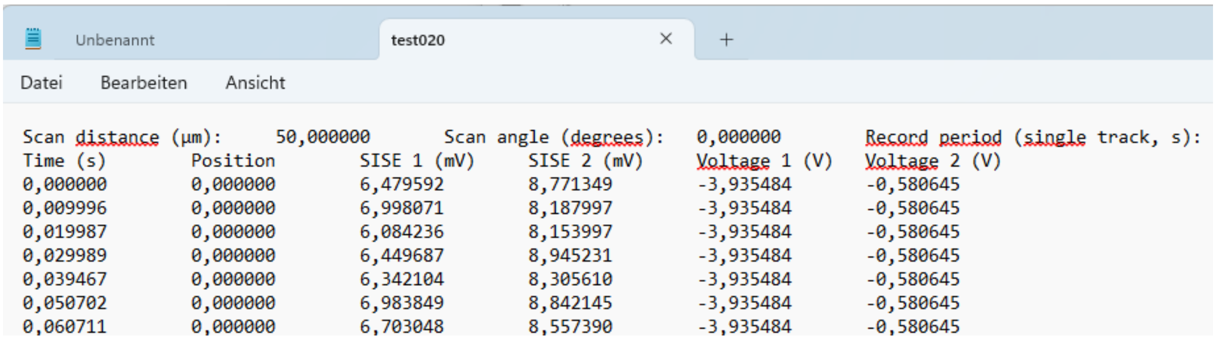


| Scan distance (µm): | 50,000000 | Scan angle (degrees): | | 0,000000 | Record period (single track, s): |
|---|---|---|---|---|---|
| Time (s) | Position | SISE 1 (mV) | SISE 2 (mV) | Voltage 1 (V) | Voltage 2 (V) |
| 0,000000 | 0,000000 | 6,479592 | 8,771349 | -3,935484 | -0,580645 |
| 0,009996 | 0,000000 | 6,998071 | 8,187997 | -3,935484 | -0,580645 |
| 0,019987 | 0,000000 | 6,084236 | 8,153997 | -3,935484 | -0,580645 |
| 0,029989 | 0,000000 | 6,449687 | 8,945231 | -3,935484 | -0,580645 |
| 0,039467 | 0,000000 | 6,342104 | 8,305610 | -3,935484 | -0,580645 |
| 0,050702 | 0,000000 | 6,983849 | 8,842145 | -3,935484 | -0,580645 |
| 0,060711 | 0,000000 | 6,703048 | 8,557390 | -3,935484 | -0,580645 |

Figure 2, SISE-Monitor data file opened in the editor of windows. The first row shows parameters that were selected during initiation of the SISE-Monitor, the second line indicators of stored data values and the lines below show data.

These files can be opened and analysed in the SISE-Analyser, which has four main modules that are meant to be executed sequentially.

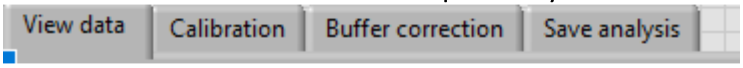


#### D. Start of the application and data browsing.

The program is started by clicking the white arrow in the upper left corner of the screen. Once activated the program can be stopped at any time by pressing the red button .

The “View data” module will get active, which has two sub-modules that can be selected in the window located in the upper left corner, named “module selection view data”. Selection will activate switches and windows of the respective sub-module.

D1. Select the “Load data” sub-module and choose the data file that should be analysed in the “Raw data file in” window. If a proper data file is chosen, the number of data points during a measurement at one position (position loop) will appear in the “Trace length” window, while the number of traces are shown in the “# of traces” window. The available SISE and Voltage channels are indicated by virtual LEDs. The three lower windows show the “Scan distance (in µm), the “Scan angle” (in °) and “Record period” (in s).

D2. If the “Scroll data” sub-module (window in upper left corner) is selected, the data can be viewed in a similar manner as during the measurement with the SISE-Monitor.

a. The switches in the middle of the “Scroll data” window allow to step, or scroll between the measurements, of which graphs are displayed in the middle window. Use the slider at the bottom of the window to adjust the speed by which data are scrolled. During scrolling, the voltage difference between position 0 and position 1 (dV) is calculated with the aid of the data that are indicated by a red “shadow” behind the white curve in the central graph. The slider “Start fit” can be used to modify the period of the measurement that is used in the dV analysis. Note that the dV values are calculated during data viewing and therefore all traces need to be viewed before the analysis can be continued. The virtual LED below “Scroll data” will turn green as soon as all data have been analysed.
In the “Event marks” window, the event marks are displayed, which were saved during the measurements.

b. The upper graph shows movement the electrode. Position 0 has the value of 0, while the movement to position 1 is shown in µm and plotted against time in seconds.

c. Voltage data of all 4 channels are plotted against time in the central graph. Each of the channels can be selected by clicking on one of the following headers;
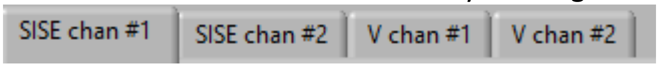

In the SISE windows, part of the white voltage trace is marked by a red “shadow”. This period of the measurement will be used for calculation of the change in voltage. The period can be attenuated with the “Start fit” slider in the lower left window.

d. In the lower graph, the procedure is depicted that is used to calculate the voltage difference (dV) between position 0 and 1. This procedure has been implicated to reduce the impact of voltage-drift of the electrode on the dV values (Newman, 2001). The procedure uses the last part of voltage measurements, as indicated by the “red shadow” in the central graph (A2c). The voltage change is determined with three measurements, in which a first regression line is calculated with the first and last data set. A second regression line, with the same angle as that of the first one, is determined for the middle data set. The voltage difference between position 0 and position 1 is calculated as the distance between both regression lines. Graphs can be selected for the SISE channel 1 and 2.

Once all data have been analysed (virtual LED below "Scroll data" is green) the header of the calibration window can be clicked to calculate ion concentrations and fluxes.

### E. Calculation of ion concentrations and fluxes

The "calibration" module is used to determine ion concentrations and -fluxes from the voltage (V) and differential voltage (dV) values, which were displayed in the "View data" window. The calibration module has 4 submodules that need to be subsequentially selected in the window "Module selection".

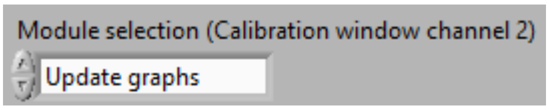


All procedures can be conducted for the SISE channels 1 and 2, which are activated by clicking the headers "SISE channel 1" or "SISE channel 2".

E1. Select the "Ion species" sub-module and choose the ion that is conducted by the ion selective electrode, values of ion mobility and valence will show up in the next two windows. The mean concentration of the measured ion (for which the electrode is selective) needs to be entered in the left window in mM (mol $m^{-3}$). The virtual LED will turn green, if an ion species is chosen and a concentration value has been entered.

E2. A calibration curve of the ion selective electrode is calculated in the sub-module "Calibration". Enter the concentration of the ion (in mM) and the voltage that was measured, for each of the calibration solutions. Data of up to 5 calibration solutions can be entered. The graph in the "Calibration data" window shows the calibration line (on a logarithmic X-scale), while the slope and intercept with the X-axis are presented above the graph. The virtual LED will turn green, when a calibration graph has been determined.

E3. In the "Geometry" submodule a correction is calculated for the scan distance of tissues (or cells) that can be regarded as cylindrical, or spheric. Once this sub-module is selected, the shape of the tissue (or cell) is chosen, by clicking one the headers: "flat surface", "cylinder", or "sphere". In case of a flat surface, the scan distance needs to be entered (in µm) and no correction occurs. Cylinders and spheres require values for the diameter of the tissue (or cell) and the minimal distance (at position 0) to the object of study. The corrected scan distance will appear in the "Corrected distance" window and the virtual LED above this window will turn green, as soon as proper values are provided.

E4. Graphs that show the calculated ion concentrations (upper graph) and ion fluxes (lower graph), plotted against time, will be generated if the "Update graphs" sub-module is chosen in the "Module selection" window. The Y-axis of both graphs can be modulated with the "Base line" and "Range limit" sliders on the left side of both graphs. The time window (X-axis) is altered with the sliders above the graphs, which affect all graphs in the sub-module.

**F. Buffer correction**

If ions that are buffered in the bath solution, the SISE measurement will underestimate the ion flux across the plasma membrane (Arif *et al.*, 1995). In buffered solutions, part of the ion flux across the plasma membrane will be compensated by a flux of the conjugated buffer, which is not taken into account for the flux calculation. With respect to pH buffers, Arif et al. (1995) developed a procedure to compensate for fluxes of conjugated ions. This procedure is implicated in the "Buffer correction" module that can be selected after the Calibration module has been completed. One of both SISE channels can be selected with the headers "SISE channel 1" or SISE channel 2" in the main "Buffer correction" window. Three sub-modules need to be activated subsequently in the "Module selection" window.

F1. Choose the buffer that was used in the "Select buffer" window, the mobility of the buffer and its $pK_S$ are shown in the windows in the middle and on the right, respectively. Some pH buffers with several acid groups (like Citrate) may be listed multiple times. Select the buffer entry of which the pH is close to the pH of the experimental solution. The virtual LED will turn green as soon as a buffer is chosen.

F2. Enter the total concentration of the buffer (conjugated + non-conjugated buffer) of the experimental solution (in mM). The virtual LED turns green if a buffer concentration is provided.

F3. In the "Update graphs" sub-module, a graph will be shown with ion flux values that are compensated for the flow of conjugated $H^+$-buffer. These values thus will represent the total flux of ions across the plasma membrane of the tissue, or cell, that is studied.

**G. Save analysis**

Results of the analysis are stored in an analysis file in the "Save analysis" module. In the "Save analysis" window, 3 sub-modules need to be addressed, which can be selected in "Module selection" window.

G1. In the "Create analysis file" sub-module a non-existing file must be entered in the "analysis output file" window. The virtual LED will turn green if a file name has been provided.

G2. The channels that are available for storage, are indicated with virtual LEDs on the left side of the table in the “Save analysis file” window. On the right side of the table data can be selected for storage, using the switch next to each parameter. The data will be stored after clicking the “Save data” switch and the virtual LED below “Save analysis file” will turn green. The data will be stored in a tab-delimited text file (Fig. 3), which can be imported into spreadsheet programs, like Excel.

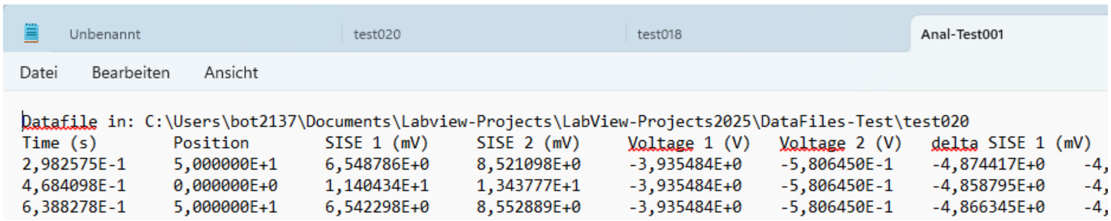


Datafile in: C:\Users\bot2137\Documents\Labview-Projects\LabView-Projects2025\DataFiles-Test\test020

| Time (s) | Position | SISE 1 (mV) | SISE 2 (mV) | Voltage 1 (V) | Voltage 2 (V) | delta SISE 1 (mV) | |
|---|---|---|---|---|---|---|---|
| 2,982575E-1 | 5,000000E+1 | 6,548786E+0 | 8,521098E+0 | -3,935484E+0 | -5,806450E-1 | -4,874417E+0 | -4, |
| 4,684098E-1 | 0,000000E+0 | 1,140434E+1 | 1,343777E+1 | -3,935484E+0 | -5,806450E-1 | -4,858795E+0 | -4, |
| 6,388278E-1 | 5,000000E+1 | 6,542298E+0 | 8,552889E+0 | -3,935484E+0 | -5,806450E-1 | -4,866345E+0 | -4, |

Figure 3, SISE-Analysis file opened in the editor of windows. The first row shows the path and filename of the data file that was analysed. The second line indicates the parameters that are stored, and the lines below show data.

G3. Switches in the “Save analysis” window allow the user to start a new data analysis or terminate the SISE-Analyser program.

### Protocol for testing the SISE-setup

The SISE-set up can be tested with a capillary that is filled with an acidic pH buffer and placed in a more alkaline bath solution. This will provide a prolonged flux of $H^+$ from the tip of the capillary that can be easily detected with a scanning pH electrode. Below the protocol and setup of this SISE test system are described.

**Preparation of the ion-selective electrodes**

Detailed protocols for preparing ion-selective electrodes that can be used to detect ion fluxes have been provided in several papers (Shabala *et al.*, 2012; Luxardi *et al.*, 2015; Portes & Feijo, 2021). Here we will summarise the steps that were executed to prepare the ion selective electrodes for our measurements.

Electrodes were pulled from borosilicate capillaries of 1.5 mm outer- and 1.05 mm inner diameter and 80/100 mm length, without filament (GB150T-8P, Science Products, Hofheim, Germany, or product Nr. 1408411, Hilgenberg, Malsfeld, Germany). The electrodes were first pulled in a two-step procedure with a vertical electrode puller (PC-10, Narishige, Tokyo, Japan). The tip size was increased to 10 - 15 µm by breaking it back against a metal wire, this procedure was monitored with a microscope. The electrodes were placed horizontally in an aluminium box (200 x 100 x 10 mm), which was heated overnight in an oven at 220° C. The next day, the capillaries were silanized by injecting 30 µl N,N-DiMethylTriMethylSilylamine, via a hole in the lid of the aluminium box. Thereafter, electrodes were kept for another 0.5 h at 220° C and subsequently cooled down.

Silanized capillaries were back filled with an electrode solution containing 10 mM KCl and 10 mM KMes pH 6.0 (Tab. 1). The tip of the capillary was viewed under a microscope and a pressure was applied at the back of the capillary to ensure that the tip of the electrode was filled with solution. The pressure was released again, just before the tip was placed in a drop of $H^+$-ionophore (Hydrogen ionophore I – Cocktail B, Sigma-Aldrich, St. Louis MO-IL, USA). Because of the hydrophobic nature of the silanized capillary the ionophore was drawn into the tip. This was continued until 30-50 µm of the tip was filled with ionophore. The electrode was kept with the tip in a solution with 10 mM KCl until use.

**$H^+$ flux measurement**

The ion selective electrode was placed in a half-cell, filled with electrode solution, and of which an Ag/AgCl wire (0.25 mm diameter) was placed in the electrode. The half-cell was connected to the headstage of micro electrode amplifier (input impedance > $10^{14}$ Ω, HS111, Bio-Logic, Claix, France). The headstage was fixed to a micromanipulator (Patchstar, Scientifica, Uckfield, UK), which was used to place it with the tip in the lid of a 90 mm Petri dish, filled with bath solution (10 mM KCl and 0.1 mM KMes pH 6.0, Tab. 1). A $H^+$ source capillary also placed in the bath, of which the tip was pulled on the PC-10 electrode puller and broken back with tweezers to 100 µm. The tip of this capillary was filled with 100 mM KCitrate pH 4.0 and 2% Agarose and the remaining of the capillary was backfilled with 100 mM KCitrate pH 4.0 (Tab. 1). A third borosilicate glass capillary (1 mm outer diameter, 0.56 mm inner diameter, 100 mm length) served as reference electrode, which was filled with 10 mM KCl (Tab. 1), plugged with 10 mM KCl in 2% agarose and placed in a half-cell with an Ag/AgCl wire (0.25 mm diameter).

Before the start of the experiment, the tip of the ion selective electrode was positioned at 50 µm distance of the tip of the $H^+$ source capillary. The electrode was moved horizontally between two positions, with a scan distance of 100 µm, and a period of 10 s.

**Table 1** Overview of solutions used in the SISE-test experiment

| Solution | Composition |
|---|---|
| Electrode | 10 mM KCl and 10 mM KMes pH 6.0 |
| $H^+$ source | 100 mM KCitrate pH 4.0 (plugged with same sol. in 2% agarose) |
| Bath | 10 mM KCl and 0.1 mM KMes pH6.0 |
| Reference | 10 mM KCl (plugged with same sol. in 2% agarose) |
| Ionophore | Hydrogen ionophore I – Cocktail B |

**Microelectrode amplifier and noise filter**

The HS111 headstage was connected to a customized version of the VF102 dual micro electrode amplifier (Bio-Logic), with a low pass filter module as shown in Fig. 1. This module amplifies the voltage signal by a factor of -33, which is corrected in the SISE-Monitor software. The output of this filter module was connected via a BNC cable, to a NI-USB-6259 analog/digital converter. In turn, the analog/digital converter was connected with a USB cable to a windows computer, running the SISE-Monitor application. Please refer to the SISE-Monitor and SISE-Analyser manual for instructions to conduct and analyse the ion flux measurements.

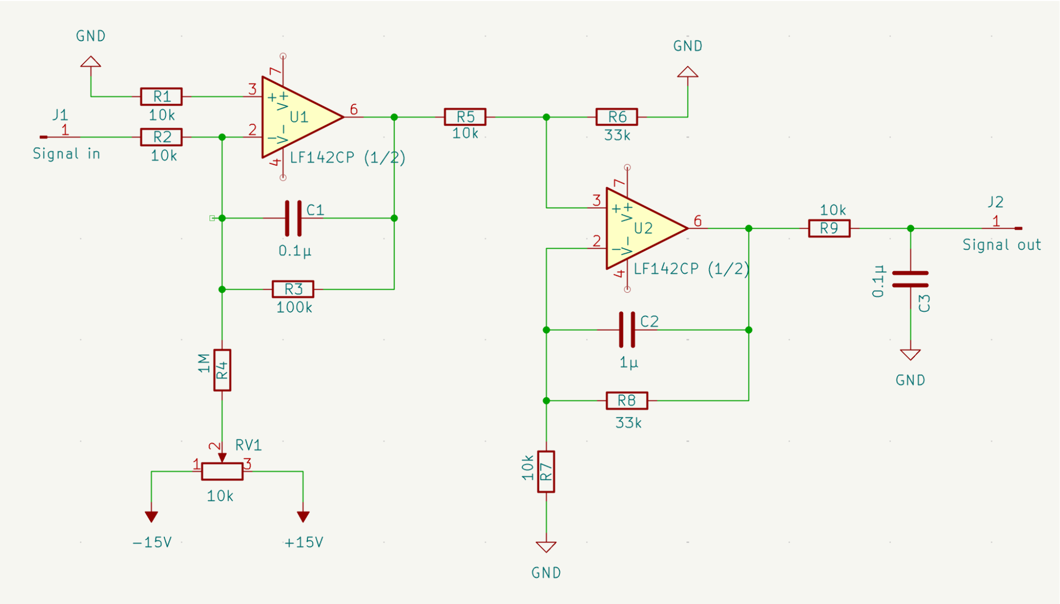


**Fig. 1** Electronic circuit of low pass active filter module mounted into the microelectrode amplifier. Circuit comprises an inverting low pass filter, followed by a non-inverting low pass filter element.

### Results

During the SISE-measurement with the $H^+$ source capillary, changes in the electrode voltage can be approximately 50 mV, as shown in Fig. 2. Note the “View data” window of the SISE-Analyser is shown in Fig. 2, in which data are shown as voltage (Panel D) and differential voltage data (Panel E). The $H^+$-concentrations and -fluxes are determined in the next “Calibration” module.

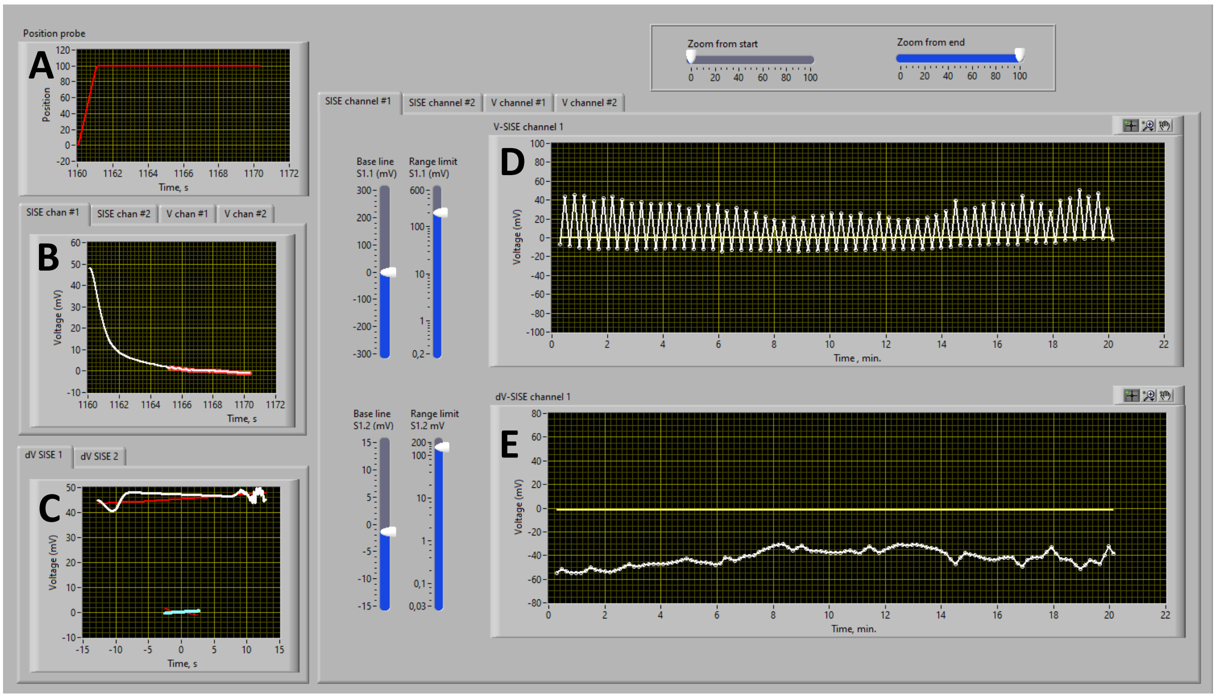


**Fig. 2.** Graphs of the SISE-Analyser obtained during evaluation a measurement of H+-fluxes from a $H^+$ source capillary, as describe above. **A.** Change in position of the electrode from position 0 (0 µm) to position 1 (100 µm), plotted against time (s). **B.** Change in voltage recorded by the ion selective electrode, plotted against time (s), the “red shadow” behind the white curve indicates that period of measurement that is used to determine average voltage and differential voltage values. **C.** Voltage difference between position 1 (measurement shown in A and B, t = -2.5 until 2.5 s) and at position 0 before (t = -13 until -8 s) and after (t = 8 until 13 s) this measurement. A first regression line (red line) was calculated for the data obtained at position 0 and a second regression line (light blue line), with the same angle as the first line, was calculated for data at position 1. **D.** Graph shows voltage signals alternating between position 1 and position 0 in time (min.). **E.** Graph shows voltage difference between position 0 and position 1, negative values represent the efflux of $H^+$.

### References protocol

**Arif I, Newman IA, Keenlyside N. 1995.** PROTON FLUX MEASUREMENTS FROM TISSUES IN BUFFERED SOLUTION. *Plant Cell and Environment* **18**(11): 1319-1324.

**Barry PH, Lynch JW. 1991.** Liquid junction potentials and small-cell effects in patch-clamp analysis. *Journal of Membrane Biology* **121**(2): 101-117.

**Luxardi G, Reid B, Ferreira F, Maillard P, Zhao M. 2015.** Measurement of Extracellular Ion Fluxes Using the Ion-selective Self-referencing Microelectrode Technique. *Jove-Journal of Visualized Experiments*(99): 11.

**Newman IA. 2001.** Ion transport in roots: measurement of fluxes using ion-selective microelectrodes to characterize transporter function. *Plant Cell and Environment* **24**(1): 1-14.

**Ng B, Barry PH. 1995.** The measurement of ionic conductivities and mobilities of certain less common organic ions needed for junction potential corrections in electrophysiology. *Journal of Neuroscience Methods* **56**(1): 37-41.

**Portes MT, Feijo JA. 2021.** Measuring Extracellular Proton and Anionic Fluxes in Arabidopsis Pollen Tubes. *Bio-protocol* **11**(3): e3908.

**Shabala S, Cuin TA, Shabala L, Newman I. 2012.** Quantifying kinetics of net ion fluxes from plant tissues by non-invasive microelectrode measuring MIFE technique. *Methods in molecular biology (Clifton, N.J.)* **913**: 119-134.

**Vanysek P 2012.** Ionic conductivity and diffusion at infinite dilution. In: Haynes WM ed. *CRC Handbook of Chemistry and Physics*. Boca Raton, Fl: CRC Press, Taylor and Francis group.

**Ion mobility data**

The $K^+$ mobility was deduced from its diffusion constant (Vanysek, 2012) using equation 1 (Newman, 2001).

Equation 01, $$u = \frac{D}{R\,T}$$

, where u is the mobility of the ion, D the diffusion coefficient, R the gas constant and T the absolute temperature. A diffusion coefficient of 1.96 $10^{-5}$ $cm^2$ $s^{-1}$ was obtained for $K^+$ (Vanysek, 2012). Based on values of R (8.3) and T (298 °K), a mobility value 7.9 $10^{-13}$ $(m.s^{-1})(N\ mol^{-1})^{-1}$ was calculated for $K^+$.

All other ion mobility values in Tab. 1 were calculated from ion mobility values relative to $K^+$, provided in Table 1 of Barry and Lynch (1991) (Barry & Lynch, 1991).

**Table 1.** Ion mobility deduced form table 1 from Barry and Lynch (1991), as explained above. These values are used in the calibration module of the SISE-Analyser to calculate ion fluxes.

| Ion species | Mobility relative to $K^+$ | Mobility ($10^{-13}$ $(m.s^{-1})(N\ mol^{-1})^{-1}$) |
|---|---|---|
| $H^+$ | 4.76 | 37,6 |
| $K^+$ | 1 | 7.9 |
| $NH_4^+$ | 1 | 7,9 |
| $Na^+$ | 0.68 | 5,4 |
| $Cl^-$ | 1,04 | 8.2 |
| $NO_3^-$ | 0.97 | 7.7 |
| $Ca^{2+}$ | 0.41 | 3.2 |
| $Mg^{2+}$ | 0.36 | 2.8 |

For buffer correction, the $pK_S$ values of the buffers were obtained from Wikipedia ($2^{nd}$ of April 2025) and the relative mobility values were acquired from Ng and Barry (1995) (Ng & Barry, 1995), or from the CRC Handbook of Chemistry and Physics (Vanysek, 2012).

**Table 2.** $pK_S$ and buffer mobility values used in the buffer correction module of the SISE-Analyser.

| Buffer species | pKS | Mobility relative to $K^+$ | Mobility ($10^{-13}$ $(m.s^{-1})(N\ mol^{-1})^{-1}$) |
|---|---|---|---|
| Mes | 6.2 | 0.37 | 2.9 |
| Malate | 5.1 | 0.40 | 3.2 |
| Citrate | 4.8 | 0.32 | 2.5 |
| TRIS | 8.2 | 0.40 | 3.2 |
| HEPES | 7.5 | 0.30 | 2.4 |
